# Characterization of vaginal microbiomes in clinician-collected bacterial vaginosis diagnosed samples

**DOI:** 10.1101/2024.10.01.616135

**Authors:** Hayden N Brochu, Qimin Zhang, Kuncheng Song, Ling Wang, Emily A Deare, Jonathan D Williams, Crystal R Icenhour, Lakshmanan K Iyer

**Affiliations:** Labcorp Center for Excellence in Data Science, AI and Bioinformatics, Burlington, NC 27215, USA; Labcorp Research and Development, Office of the CSO, Burlington, NC 27215, USA

## Abstract

Bacterial vaginosis (BV) is a type of vaginal inflammation caused by bacterial overgrowth, upsetting the healthy microbiome of the vagina. Existing clinical testing for BV is primarily based upon physical and microscopic examination of vaginal secretions, while more modern PCR-based clinical tests target panels of BV-associated microbes, such as the Labcorp NuSwab® test.

Remnant clinician-collected NuSwab® vaginal swabs underwent DNA extraction and 16S V3-V4 rRNA gene sequencing to profile microbes in addition to those included in the Labcorp NuSwab® test. Community State Types (CSTs) were determined using the most abundant taxon detected in each sample. PCR results for NuSwab® panel microbial targets were compared against the corresponding microbiome profiles. Metabolic pathway abundances were characterized via metagenomic prediction from amplicon sequence variants (ASVs).

Sequencing of 75 remnant vaginal swabs yielded 492 unique 16S V3-V4 ASVs, identifying 83 unique genera. NuSwab® assay microbe quantification was strongly concordant with quantification by sequencing (p << 0.01). Samples in CST-I (18 of 18, 100%), CST-II (3 of 3, 100%), CST-III (15 of 17, 88%), and CST-V (1 of 1, 100%) were largely categorized as BV-negative via the NuSwab® panel, while most CST-IV samples (28 of 36, 78%) were BV-positive or BV-indeterminate. BV-associated microbial and predicted metabolic signatures were shared across multiple CSTs.

These findings show that 16S V3-V4 rRNA gene sequencing robustly reproduces PCR-based BV diagnostic testing results, accurately discriminates vaginal microbiome CSTs dominated by distinct *Lactobacilli*, and further elucidates BV-associated bacterial and metabolic signatures.

**Importance:** Bacterial vaginosis (BV) poses a significant health burden for women during reproductive years and onward. Current BV diagnostics rely on either physical and microscopic evaluations by technicians or panels of select microbes. Here, we sequenced the microbiome profiles of samples previously diagnosed by the Labcorp NuSwab® test to better understand disruptions to the vaginal microbiome during BV. We show that microbial sequencing can reproduce targeted panel diagnostic results, while also broadly characterizing healthy and BV-associated microbial and metabolic biomarkers. This work highlights a robust, agnostic BV classification scheme with potential for future development of sequencing-based BV diagnostic tools.

## Introduction

Women’s health is an issue of global concern that requires substantial effort to resolve the gender inequities in funding and quality of care that persist today^1,2^. Organizations, such as the World Health Organization (WHO), have recognized the importance of women’s health and is investing considerable resources and efforts to enhance access to healthcare for women^3^. Among the health challenges faced by women, bacterial vaginosis (BV) poses a significant health burden with a prevalence of 21.2 million women aged 14-49 as of 2004, occurring initially during reproductive years and continuing through life^4^. Healthy vaginal microbiomes can contain dozens of microbial species in a specific balance^5^, while BV is a type of vaginal inflammation caused by bacterial overgrowth that upsets the healthy microbiome of the vagina^6^.

Existing clinical testing for BV is primarily based upon physical and microscopic examination of vaginal secretions, such as Amsel’s criteria and Nugent score^7^. In Amsel’s criteria, the presence of at least three out of the following four criteria is indicative of BV: high vaginal pH (>4.5), thin and homogenous discharge, malodorous fishy discharge upon adding 10% potassium hydroxide, and identification of clue cells. The Nugent score is a scoring system that evaluates the presence of *Lactobacillus spp.* morphotypes, *Gardnerella vaginalis* morphotypes, and *Mobiluncus spp.* morphotypes. Newer nucleic acid-based clinical tests are now available. The Labcorp NuSwab® test^8,9^ is a highly sensitive and specific PCR-based method that diagnoses BV in symptomatic women by quantifying three key BV-associated microbes: *Atopobium vaginae*, *Bacterial Vaginosis Associated Bacterium (BVAB)-2*, and *Megasphaera-1*. While these diagnostics provide widely accessible clinical assessment of BV, many symptomatic women receive negative test results (roughly 50% in some studies) and have limited healthcare alternatives for further diagnostic evaluation and treatment. Some of these women test positive on broader panel vaginitis tests (e.g. NuSwab® Vaginitis Plus) that also check for other common sources of vaginal inflammation, such as *Candida spp.* and *Trichomonas vaginalis*. However, many women fail to receive diagnosis and often face negative impacts on their quality of life and mental health^10^, so it is imperative to expand and improve BV diagnostic tools available to clinicians.

16S rRNA gene sequencing, an amplicon-based next generation sequencing approach, is a more comprehensive method for characterizing the composition and diversity of vaginal microbiomes. Such sequencing-based analyses have drastically improved our understanding of vaginal microbiome diversity and the shifts in microbiome composition throughout a woman’s premenarchal and reproductive years^11–14^. Amplicon sequencing-based profiling has also revealed unique community state types (CSTs) within the vaginal microbiome, which are defined by the key *Lactobacillus* species detected^12^. *Lactobacilli* are essential for vaginal microbiome health, interacting with mucosal immune and epithelial cells via immunomodulatory mechanisms^15,16^. Disruptions to this *Lactobacillus*-mediated vaginal homeostasis have been associated with dysbiotic conditions, namely BV and aerobic vaginitis^17^. Many amplicon sequencing-based investigations of vaginal microbiomes have identified additional correlates of BV positivity, such as *Gardnerella* and *Prevotella*^18–20^. Despite these advances in our understanding of BV-associated vaginal microbiota, testing of clinical samples using PCR-based diagnostics provides limited clinical utility^21^.

In this study, we characterized the vaginal microbiomes of 75 clinician-collected remnant NuSwab® vaginal swabs via 16S V3-V4 rRNA gene sequencing. We elucidated the rich diversity and CSTs of these vaginal microbiomes with amplicon sequence variant (ASV) resolution, showing that BV typically manifests in *Lactobacillus*-deficient states. We also showed that amplicon-based sequencing accurately identifies the three microbes targeted by the NuSwab® BV PCR assay, as well as identifying additional BV-associated microbes. Using metabolic predictions from these ASVs, we identified metabolic signatures significantly associated with BV positivity across multiple CSTs. These findings show that amplicon sequencing can accurately reproduce PCR-based testing results and provide an expanded array of biomarkers that may enhance currently available diagnostic tests.

## Results

### 16S rRNA gene sequencing accurately reproduces PCR-based BV diagnostic testing results

75 remnant clinician-collected vaginal swabs were previously analyzed via the Labcorp NuSwab® test^8^, a PCR-based diagnostic method that detects bacterial vaginosis (BV) by scoring three key BV-associated microbes: *Atopobium vaginae*, *BVAB-2*, and *Megasphaera-1* (**Figure 1a**). PCR analysis of these vaginal swabs yielded BV-positive (BV-POS, 27 of 75, 36%), BV-indeterminate (BV-IND, 3 of 75, 4%), and BV-negative (BV-NEG, 45 of 75, 60%) diagnostic results. To further characterize these vaginal swabs, we profiled their microbiomes via 16S V3-V4 rRNA gene sequencing analysis. Sequencing yielded 492 unique 16S V3-V4 Amplicon Sequence Variants (ASVs) that mapped to 83 unique genera, resulting in a mean of 93.6K genus mapped reads per sample. All three NuSwab®-tested microbes were detected among these ASVs, with significant enrichment of 16S relative abundances (RAs) among samples with scores of 2 (**Figure 1b**, all p-values << 0.001, Wilcoxon rank-sum tests). This strong corroboration of BV diagnostic results by 16S sequencing analysis highlights the robustness of the NuSwab® test.

**Figure 1.**
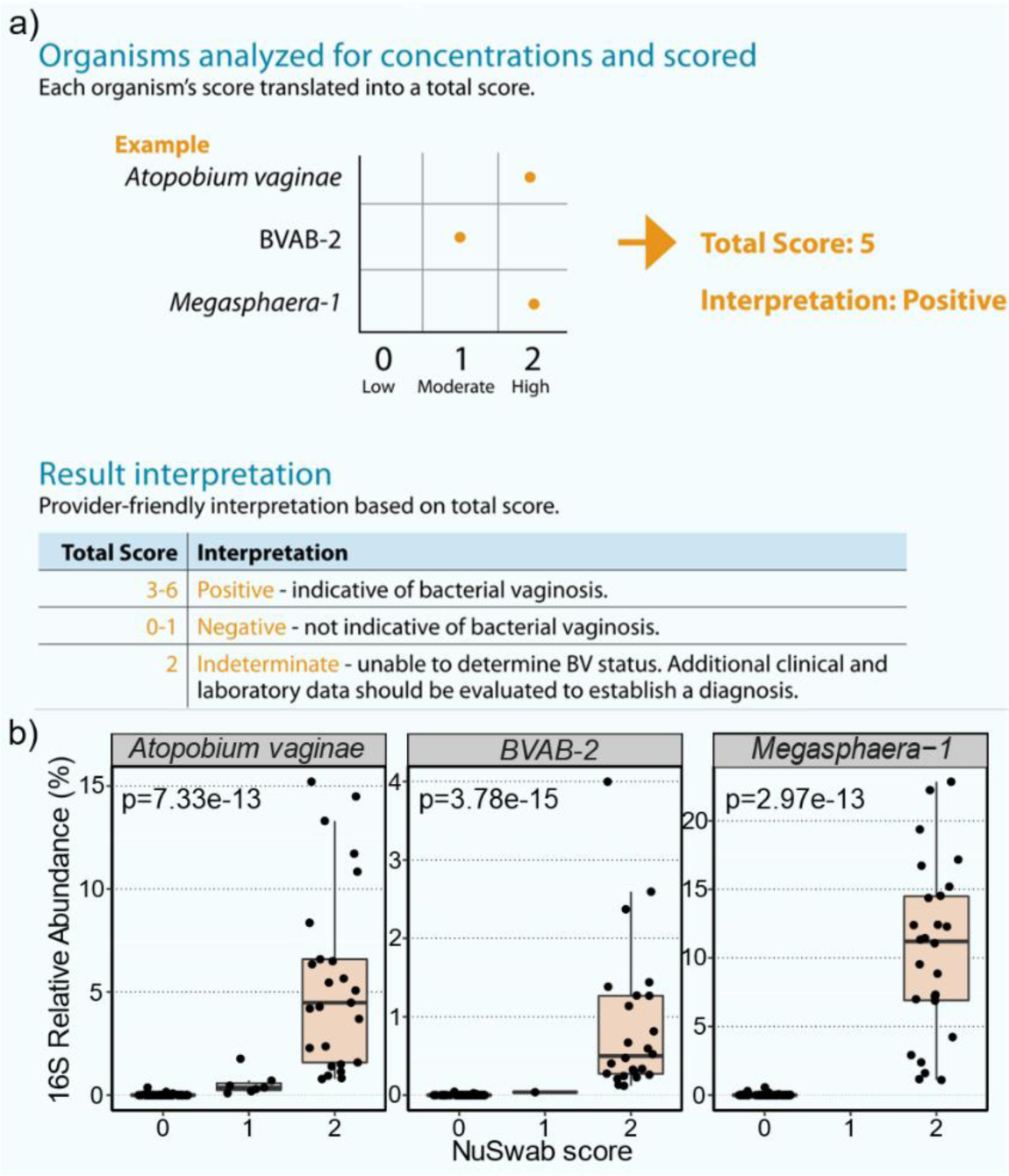
16S V3-V4 rRNA sequencing of remnant clinician-collected vaginal swabs previously analyzed via Labcorp NuSwab® BV PCR test. a) Example process of determining Bacterial Vaginosis (BV) status of a vaginal swab using the NuSwab® three microbe panel composed of *Atopobium vaginae*, *BVAB-2*, and *Megasphaera-1*. Each microbe is quantified and given a score of 0 (low), 1 (moderate), or 2 (high). The total score is then interpreted as BV-POS (3-6), BV-IND (2), or BV-NEG (0-1). b) 16S relative abundances of NuSwab® panel microbes stratified by their scores. Statistical comparisons of 16S relative abundances were made between samples with NuSwab® scores of 2 and those with scores of 0 or 1 using Wilcoxon rank-sum tests with FDR control.

We next sought to identify additional characteristics of these vaginal swabs by investigating their ASV-resolved microbiomes. Juxtaposition of samples by BV status using class-level taxonomy revealed a strong propensity for bacilli detection among BV-NEG samples (80.6% mean RA) compared to BV-IND (31%) and BV-POS (11.5%) samples (**Figure 2a**). Of the other 10 bacterial classes identified, six had an average RA of 5% or greater in BV-POS and BV-IND samples, while only one of those classes (Actinobacteria) was detected at similar levels in BV-NEG samples (**Figure 2a**). This was expected, as healthy vaginal microbiomes are known to be heavily populated with *Lactobacilli* (within Bacilli class), while BV microbiomes tend to be more diverse^22^. Phylogenetic analysis of ASVs confirmed the robustness of these taxonomic classifications, with ASVs closely clustering with those assigned to the same class (**Figure 2b**). Bacilli ASVs, consistent with their strong prevalence, constituted the plurality of ASVs (97 of 492, ∼20%) in the phylogeny and formed one large cluster of those predominantly in the *Lactobacillus* genus, while two other smaller clusters formed mostly comprising *Ureaplasma*, *Mycoplasma*, and *Bulleidia* (**Figure 2b-c**). A diverse population of 51 *Lactobacillus* ASVs was detected, most of which were speciated (**Figure 2c**, 38 of 51, 75%). In total, 13 *L. iners*, 11 *L. crispatus*, 7 *L. jensenii*, 5 *L. gasseri*, and 2 *L. hominis* ASVs were found, each forming close subgroups in the phylogeny (**Figure 2c**). These ASV-resolved, deeply sequenced vaginal microbiomes enable broad assessment of healthy and BV microbiome characteristics, expanding on BV diagnostic insights.

**Figure 2.**
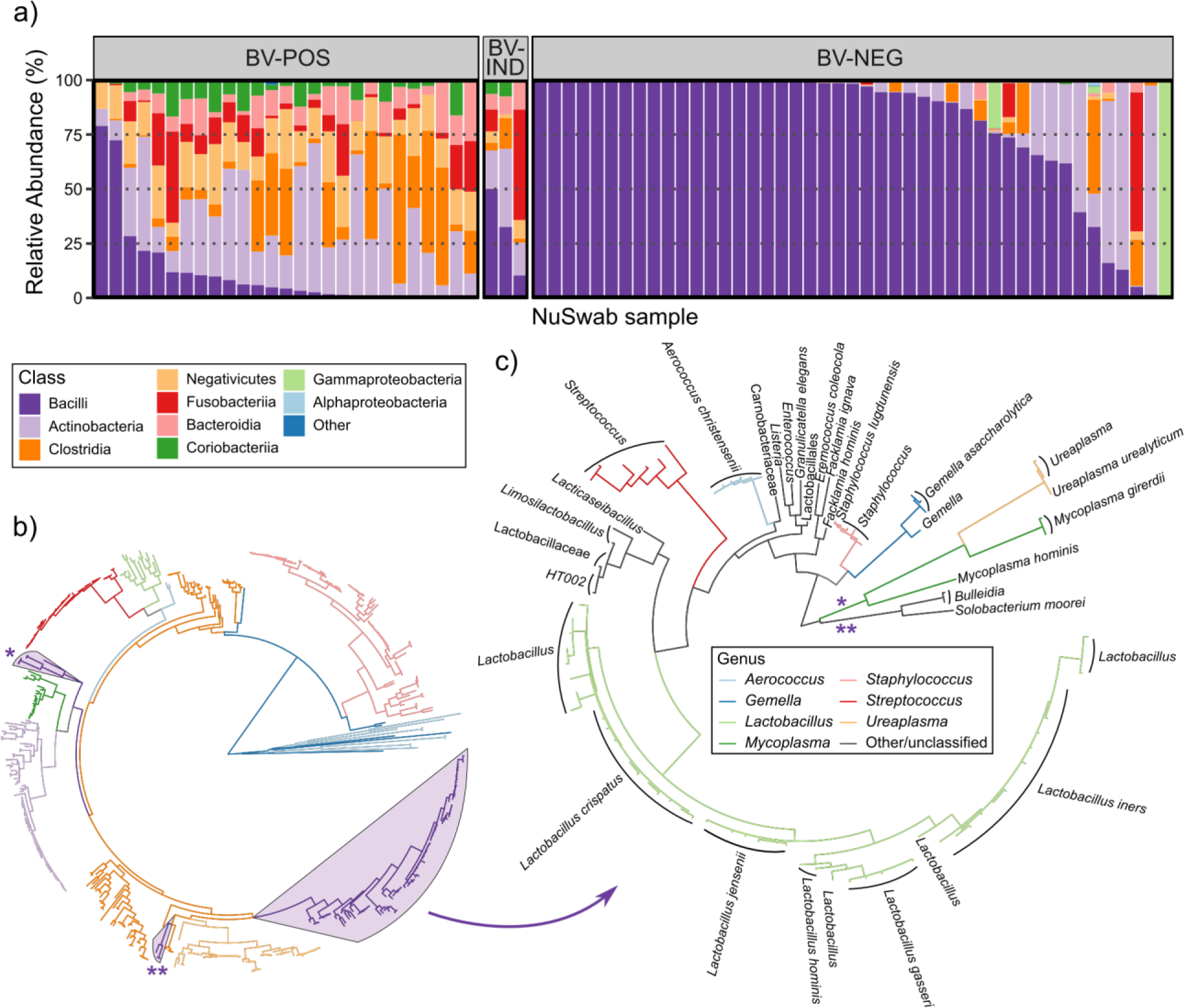
Taxonomic and phylogenetic analysis of 16S V3-V4 Amplicon Sequence Variants (ASVs). **a)** Stacked bar plot of class relative abundances colored consistently with (b) and with samples stratified by BV status. **b)** ML phylogeny of all ASVs, with branches colored by class. The Bacilli class was highlighted and constructed a phylogeny. **c)** ML phylogeny of ASVs within the Bacilli class shown in (b) with branches colored by genus and tips labeled with the lowest taxonomic classifications. Neighboring tips with the same label were aggregated into a single label.

### Characterization of community state types

Next, we further characterized these samples using community state type (CST) analysis, which differentiates samples by their detection of key *Lactobacillus* species. Specifically, we employed CST analysis per Ravel *et al*^12^, wherein five CSTs are considered based on the most abundant species detected in each sample (CST-I: *L. crispatus*, CST-II: *L. gasseri*, CST-III: *L. iners*, CST-IV: diverse communities, CST-V: *L. jensenii*). All five CSTs were observed among the 75 NuSwab® samples in this study with a strongly significant association between CST and BV status (Fisher’s exact test, p < 0.001) (**Table 1**). While most BV-NEG samples were in *Lactobacillus* dominated CSTs (37 of 45, 82%), BV-POS and BV-IND samples were largely classified in CST-IV (28 of 30, 93%) (**Table 1**, **Figure 3a**). When clustering sample microbiome profiles using multi-dimensional scaling of their pairwise Bray-Curtis Dissimilarities, we observed significant separation of samples by their CST classification (**Figure 3b**, PERMANOVA, p < 0.001).

**Figure 3.**
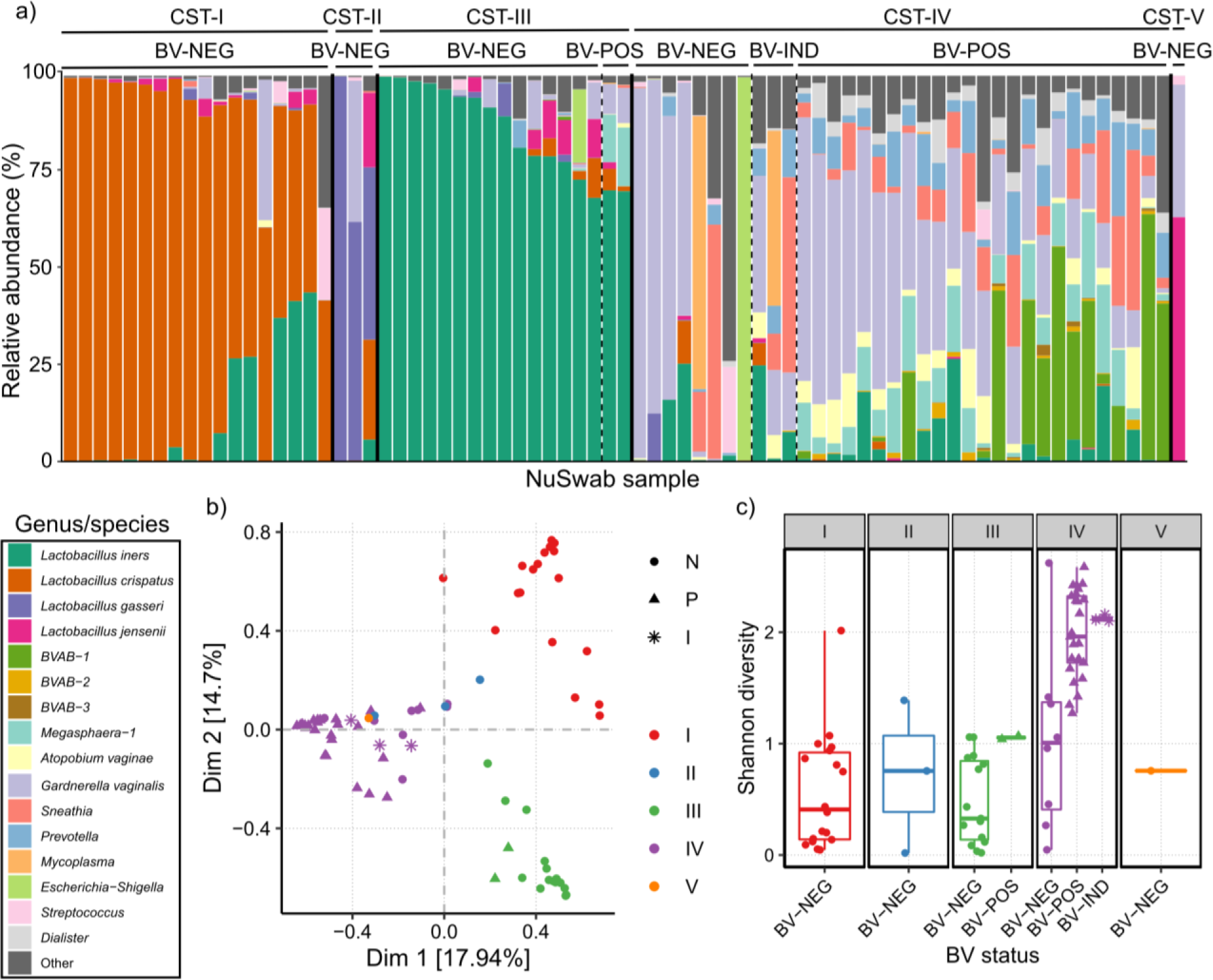
Community state type (CST) analysis of vaginal microbiomes using CST definitions described by Ravel, *et al*^12^. **a)** Stacked bar plot of key genera and species relative abundances across CSTs I, II, III, IV, and V. Samples are stratified by CST (solid vertical lines), then by BV status (dashed vertical lines). **b)** Multi-dimensional scaling scatter plot of vaginal microbiome Bray-Curtis Dissimilarities with data points colored by CST and shaped by BV status. **c)** Boxplots of vaginal microbiome diversities as measured by Shannon index, stratified first by CST, then by BV status with data points shaped and colored consistently with those in (b).

**Table 1.** Community State Type (CST) classifications of NuSwab® vaginal samples. Each table entry indicates the number of samples classified in each CST from each of the NuSwab® result categories (BV-NEG, BV-IND, BV-POS). The marginal totals are shown on the right and bottom of the table.

| NuSwab® result | CST-I | CST-II | CST-III | CST-IV | CST-V | Total |
| --- | --- | --- | --- | --- | --- | --- |
| BV-NEG | 18 | 3 | 15 | 8 | 1 | 45 |
| BV-IND | 0 | 0 | 0 | 3 | 0 | 3 |
| BV-POS | 0 | 0 | 2 | 25 | 0 | 27 |
| Total | 18 | 3 | 17 | 36 | 1 | 75 |

We also observed that BV-POS samples were notably more diverse than BV-NEG samples (**Figure 3c**), consistent with their tendency to be within CST-IV (**Table 1**, **Figure 3a**). Interestingly, within CST-IV we still observed significantly higher microbiome diversity among BV-POS and BV-IND samples compared to BV-NEG samples (**Figure 3c**, p < 0.001, Wilcoxon rank-sum test). This suggests that vaginal microbiome classification in CST-IV may represent other forms of dysbiosis not necessarily caused by BV. Together these findings indicate that among these NuSwab® samples, BV is associated with diverse, heterogenous microbiomes that generally lack the essential vaginal *Lactobacilli*.

Some studies argue that the V3-V4 hypervariable region might insufficiently resolve key vaginal *Lactobacillus* species^23,24^ (e.g. *L. iners*); therefore, we confirmed the accuracy of our CST classifications using shotgun metagenomic sequencing (MGx), which has been used to investigate species-refined vaginal microbiome diversity at the population level^25^. A separate group of 54 remnant NuSwab® vaginal samples were sequenced via both 16S V3-V4 rRNA gene sequencing and MGx, and CST results were generated using the same ruleset as above for both sequencing methods (**Table 2**). 53 of 54 (98%) samples had concordant CST calls across sequencing methods spanning all five CSTs. Closer inspection of the discrepant CST classification revealed only a difference in *L. crispatus* (16S: 37.1%, MGX: 61.4%) and *L. iners* (16S: 52.8%, MGx: 38.2%) relative abundances. Since *L. crispatus* and *L. iners* are phylogenetically distinct (**Figure 2c**), this relative abundance difference was likely due to biases from the sequencing methods rather than 16S misclassification. These results show that our 16S V3-V4 vaginal *Lactobacillus* speciation is robust and capable of yielding accurate CST classifications for vaginal swabs.

**Table 2.** Comparison of Community State Type (CST) classifications between 16S rRNA gene V3-V4 sequencing and shotgun metagenomic sequencing (MGx). Rows and columns indicate CST calls determined from MGx and 16S rRNA gene V3-V4 sequencing results, respectively.

| MGx | 16S rRNA gene V3-V4 |  |  |  |  |
| --- | --- | --- | --- | --- | --- |
|  | CST-I | CST-II | CST-III | CST-IV | CST-V |
| CST-I | 16 | 0 | 1 | 0 | 0 |
| CST-II | 0 | 1 | 0 | 0 | 0 |
| CST-III | 0 | 0 | 17 | 0 | 0 |
| CST-IV | 0 | 0 | 0 | 14 | 0 |
| CST-V | 0 | 0 | 0 | 0 | 5 |

### BV-POS samples are enriched with distinct bacterial networks

Since vaginal microbiomes exhibited clear differences in CST classifications and microbial diversity based on BV status (**Figure 3**, **Table 1**), we systematically analyzed microbial differential abundances between BV-POS and BV-NEG samples. Not surprisingly, we found all three key NuSwab® BV-associated microbes significantly enriched and multiple *Lactobacilli* species significantly depleted among BV-POS samples (**Figure 4a**). We also detected multiple other known BV-associated microbes enriched in BV-POS samples, including *BVAB-1*^26^, *BVAB-3*^26^, *Megasphaera-2*^27^, *Sneathia*^28^*, DNF00809*^27^*, and Parvimonas*^29^ (**Figure 4a-b**).

**Figure 4.**
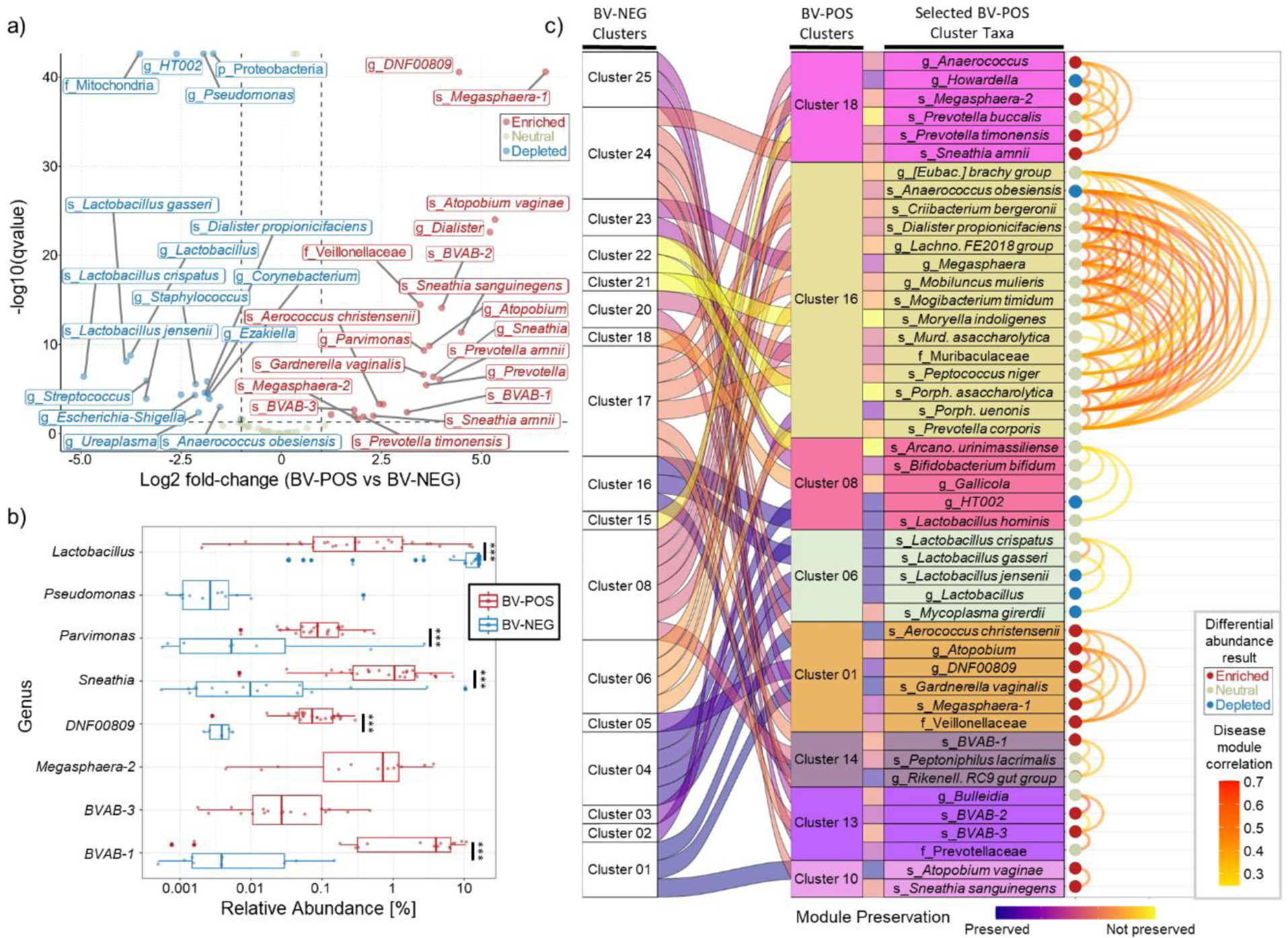
Differential abundance (DA) analysis and modularized co-occurrence network analysis of BV-POS and BV-NEG samples. **a)** Volcano plot of the differential abundance results, depicting the log2 fold-change (L2FC) (x-axis, BV-POS vs BV-NEG), and the -log10(p-value) (y-axis) for each taxa assessed. FDR control for multiple testing was used to calculate p-values. Data point colors represent statistical significance of taxa, with red representing enriched taxa (p-value < 0.05, L2FC >= 1), blue representing depleted taxa (p-value < 0.05, L2FC <= −1), and yellow representing all other taxa without significant changes. **b)** Boxplots comparing the relative abundances of select taxa between BV-POS and BV-NEG samples with statistical significance indicated to the right (*** p<0.001). Unique identification of taxa is represented by only a single boxplot shown (e.g. *Parvimonas* exclusively detected in BV-NEG samples). **c)** Clustering results from C3NA^38^ with the first column representing clustering among the BV-NEG samples (n=25 clusters) followed by the modular preservation ribbon linked to the colored BV-POS clusters (n=19). The arc on the right represents the spearman correlations above 0.2 among each BV-POS cluster, with each node colored by the DA result (enriched, neutral, depleted) and the edge colored by the correlation magnitude. All testing was performed on the species level with any non-speciated taxa labeled at the highest resolved taxonomic levels, i.e., “g_*DNF000809*” represents the ASVs that resolved to the *DNF000809* genus without species assignment.

We also evaluated the modularity of these microbes using a correlation-based analysis that assessed microbe-microbe correlations separately for BV-POS and BV-NEG samples (Methods). This approach enabled us to simultaneously evaluate correlation network conservation and whether differentially abundant microbes coordinate with each other based on BV status. Interestingly, each NuSwab® BV-associated microbe was found in a separate correlation-based cluster, suggesting that they broadly capture BV microbial signatures (**Figure 4c**). *Atopobium vaginae* formed a small cluster with only *Sneathia sanguinegens* (Cluster 10), while *BVAB-2* clustered with *BVAB-3* (Cluster 13) and *Megasphaera-1* formed a broad cluster (Cluster 01) with multiple microbes enriched in BV-POS samples (**Figure 4c**). We also identified a fourth cluster of BV-associated microbes (Cluster 18), including *Anaerococcus*, *Megasphaera-2*, *Prevotella timonensis*, and *Sneathia amnii* (**Figure 4c**). These results indicate that multiple coordinated microbial networks may drive BV pathogenesis and further support the use of multiple diagnostic targets for BV detection.

### Prediction of a BV-associated metabolic signature

To better understand the functional capacity of BV microbial signatures, we determined the metagenomic predictions of ASVs and performed differential abundance analysis of the MetaCyc^30^ metabolic pathways detected using PICRUSt2^31^. Since only few samples were classified as CST-II and CST-V (**Table 1**), we constrained this analysis to CST-I, CST-III, and CST-IV, and investigated whether metabolic pathways were significantly enriched or depleted (FDR < 0.05) based on BV positivity within and between CSTs. Since all but two BV-POS samples were in CST-IV, only CST-IV BV-POS samples were considered. Comparisons of CST-I and CST-III BV-NEG samples versus the BV-POS samples separately revealed 270 and 232 differentially abundant (DA) metabolic pathways, respectively (**Table 3**, **Supplementary Data**). When comparing the CST-IV BV-NEG samples, 36 DA metabolic pathways were detected, all of which were detected in the other two comparisons (**Table 3**, **Supplementary Data**). Furthermore, a separate comparison of BV-NEG samples between CST-I and CST-III revealed only 39 DA metabolic pathways, which only represented a small fraction of the pathways in the other comparisons (**Table 3**, **Supplementary Data**). This indicated that the BV-associated metabolic pathways were largely not due to a deficiency of *Lactobacilli* and were more likely driven by the microbes enriched in BV-POS samples (**Figure 4**).

**Table 3.** Differential abundance analysis of predicted metabolic pathways between BV-NEG and BV-POS samples. Each row shows the number of significantly differentially abundant (DA, FDR < 0.05) metabolic pathways between BV-NEG and BV-POS samples within and between CSTs. In all comparisons, the BV-POS samples were from CST-IV, hence only the BV-NEG CST is shown in the table. The total number of DA pathways is shown on the left followed by the number of pathways that were also DA between BV-NEG samples compared across CST-I and CST-III. On the right, the remaining pathways are shown with the total followed by the number shared with other comparisons and the number unique to the comparison.

| BV-NEG | Total | CST-I BV-NEG vs.<br>CST-III BV-NEG<br>(39 total) | BV-NEG vs. BV-POS |  |  |
| --- | --- | --- | --- | --- | --- |
|  |  |  | Total | Shared | Unique |
| CST-I | 270 | 36 | 234 | 194 | 40 |
| CST-III | 232 | 30 | 202 | 194 | 8 |
| CST-IV | 36 | 3 | 33 | 33 | 0 |

To visually examine the conservation of the four sample comparisons made above (**Table 3**, including the BV-NEG CST-I versus CST-III comparison), we selected the most variable DA pathways from each comparison and merged them into a single heatmap (**Figure 5**). Consistent with our DA results (**Table 3**, **Supplementary Data**), BV-POS samples showed highly similar patterns of enriched and depleted pathways regardless of CST (**Figure 5**). Interestingly, the three BV-NEG samples within CST-IV (the three left-most samples within BV-NEG CST-IV in **Figure 5**) exhibit a similar pattern of pathways with BV-POS samples. This similarity suggests a potential transition from a healthy state to BV, offering a plausible explanation for some patients experiencing BV symptoms while testing negative. Among the pathways significantly enriched in BV-POS samples in all comparisons with BV-NEG samples, we observed four noteworthy pathways: acetyl-CoA fermentation to butanoate, L-glutamate and L-glutamine biosynthesis, succinate fermentation to butanoate, and pyruvate fermentation to propanoate. These enrichments were indicative of higher concentrations of four known BV-related metabolites, namely acetate, succinate, butanoate, and propanoate^32,33^. Overall, these findings indicate that BV induces large metabolic disruptions in the vaginal microenvironment.

**Figure 5.**
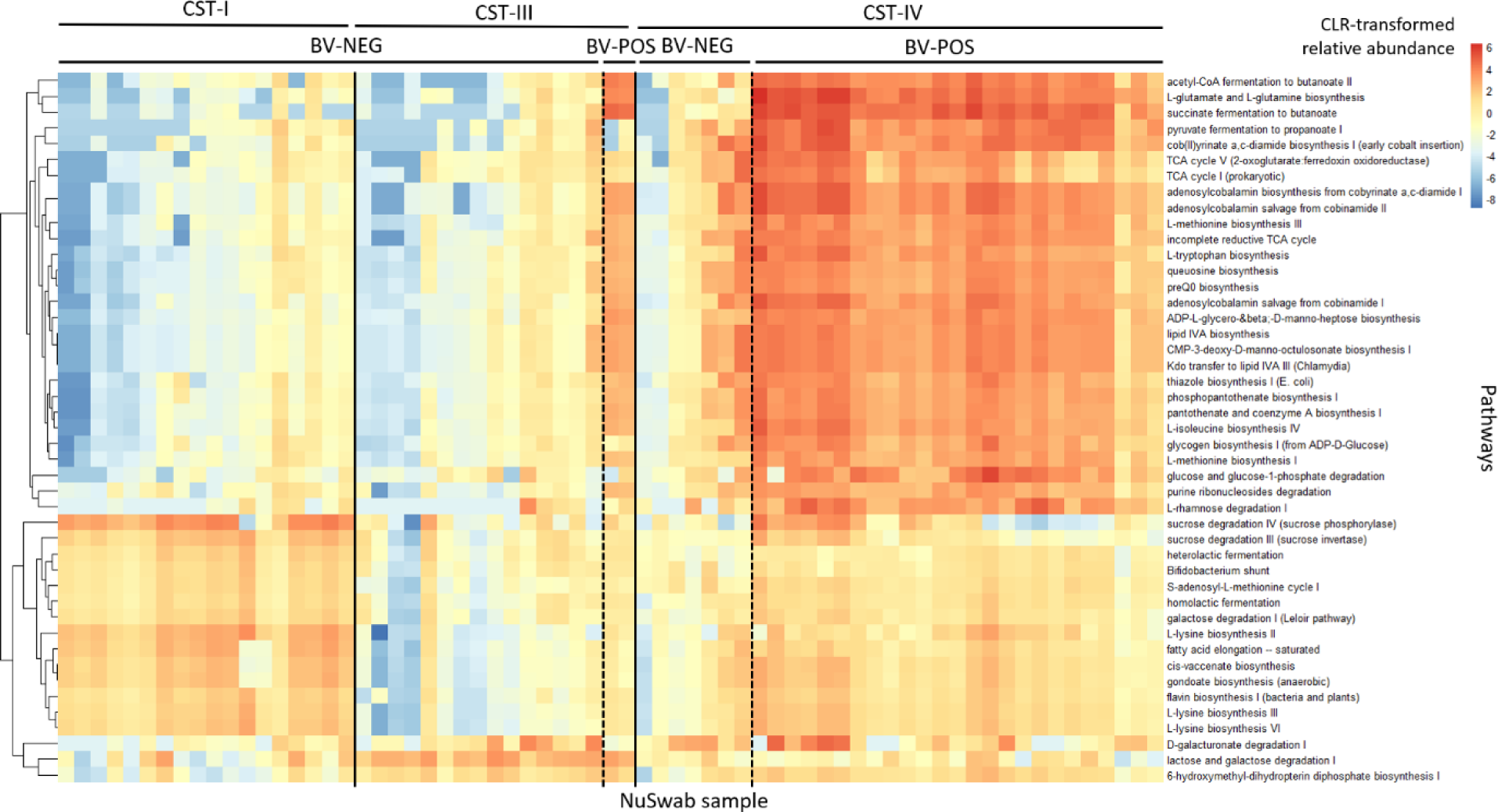
Heatmap of differentially abundant (DA) predicted metabolic pathways between BV-POS and BV-NEG samples found in CST-I, CST-III, and CST-IV. Rows represent MetaCyc^30^ metabolic pathways and are ordered via hierarchical clustering, while columns (samples) are stratified first by CST (solid vertical lines) then by BV status (dashed vertical lines). The heatmap combines the results of four pairwise comparisons, including: 1) CST-I BV-NEG vs CST-III BV-NEG, 2) CST-I BV-NEG vs CST-IV BV-POS, 3) CST-III BV-NEG vs CST-IV BV-POS, and 4) CST-IV BV-POS vs CST-IV BV-NEG. The top 20 differentially abundant pathways of each comparison with the greatest variance were selected and merged into the list of significant pathways across all pairs of sample groups (combined set of 45 pathways shown).

## Discussion

In this study, we comprehensively profiled the microbial and metabolic signatures of remnant clinician-collected, PCR tested healthy and BV sample microbiomes. We demonstrated that 16S V3-V4 rRNA gene sequencing can 1) reproduce Labcorp NuSwab® BV PCR testing results; 2) accurately classify vaginal microbiome CSTs via *Lactobacillus* speciation; and 3) identify additional microbial and metabolic correlates of BV positivity.

By comprehensively profiling the vaginal microbiome with ASV-level resolution, we elucidated multiple features of BV positivity, confirming many previous findings. Not surprisingly, we found that BV-POS samples tended to be deficient in *Lactobacillus* species relative to BV-NEG samples, and instead were significantly more diverse and enriched with multiple BV-associated bacteria, such as *Gardnerella*, *Prevotella*, *Sneathia*, *Dialister*, *BVAB-1*, *BVAB-3, veillonellaceae*, *DNF00809*, *Aerococcus christensenii, Parvimonas,* and *Bulleidia*^26,27,34–37^. The three NuSwab® BV-associated microbes (*Atopobium vaginae*, *BVAB-2*, and *Megasphaera-1*) were also among the enriched bacteria in BV-POS samples. Interestingly, we observed weak enrichment of *Gemella asaccharolytic*a in BV-POS samples, whereas *Gemella spp.* has been previously found to be less frequently detected in BV-POS samples^26^. We also uniquely evaluated the correlation networks of BV-associated bacteria using C3NA^38^, highlighting key modular differences between the BV-POS and BV-NEG samples in this study. We uncovered four clusters of BV-associated microbes, and intriguingly the three NuSwab® BV-associated microbes were identified in three separate clusters, indicating that those microbes collectively capture a significant portion of BV microbial signatures. One of those clusters was comprised of *Gardnerella vaginalis*, *Megasphaera-1*, *DNF00809*, and several other taxa, which could represent a group of BV biomarkers worth investigating as alternative therapeutic treatments or to enhance current diagnostic testing capabilities. However, while *Gardnerella vaginalis* was enriched among BV-POS samples in this study and is targeted by some commercially available PCR tests, it is important to note that *Gardnerella vaginalis* is also prevalent among women who do not have BV^39^. Evaluation of BV positive samples from multiple population cohorts might be necessary to critically assess the diagnostic utility of these microbial biomarkers.

In this study we also interrogated the metabolic signatures that distinguish vaginal microbiomes based on their CST and/or BV diagnostic status. We observed four noteworthy pathways: acetyl-CoA fermentation to butanoate, L-glutamate and L-glutamine biosynthesis, succinate fermentation to butanoate, pyruvate fermentation to propanoate. These enrichments indicated higher concentrations of four known BV-related metabolites, namely acetate, succinate, butanoate, and propanoate^32,33^. Acetate and butanoate are known to play a role in immune responses^40^. We also observed three pathways significantly depleted within CST-I, namely D-galacturonate degradation, lactose and galactose degradation and 6-hydroxymethyl-dihydropterin diphosphate biosynthesis, and some pathways significantly depleted in CST-III, such as sucrose degradation and L-lysine biosynthesis. The observed depletion of L-lysine biosynthesis in CST-III corresponds with findings from prior studies, which have indicated that *L. iners*, the dominant species in CST-III, has a considerably smaller core genome for the biosynthesis of lysine compared to *L. crispatus*, the dominant species in CST-I^41^. These metabolomic insights represent a complementary suite of biomarkers indirectly detectable by 16S rRNA gene sequencing and suggest that metabolomic studies could elucidate immune dysfunction in the vaginal microenvironment during BV pathogenesis.

Here, we demonstrated that 16S rRNA gene sequencing provides robust detection of BV-associated microbial and metabolic signatures. However, by only targeting certain hypervariable regions (V3-V4 in our case), there are limitations to the resolution, and this may vary by genus. Fortunately, we show that the V3-V4 hypervariable regions accurately speciate key strains of *Lactobacillus*, but other genera, such as *Gardnerella*, had fewer species resolved. A recent study using shotgun metagenomic sequencing of vaginal microbiomes showed differences in *Gardnerella* species richness across multiple cohorts of pregnant women^42^, highlighting the importance of its species-level resolution. Thus, limitations in 16S rRNA amplicon taxonomic resolution could limit its use in the clinic. However, full-length 16S rRNA gene sequencing has been shown to improve taxonomic resolution and resolve single nucleotide differences in the entire 16S rRNA gene^43^. In practice, full-length 16S rRNA gene sequencing has already been shown to significantly improve species-level accuracy using more than one sequencing platform, highlighting its potential clinical utility^44–47^. Moreover, a recent study showcased a bioinformatics workflow tailored for long-read 16S rRNA gene sequencing of vaginal microbiomes suitable in a clinical setting^48^. Shotgun metagenomic and metatranscriptomic sequencing have also provided highly resolved insights into the vaginal microenvironment^49^, including fungi, parasites, and DNA viruses, suggesting potential for clinical diagnostics as well. However, the relatively lower cost of amplicon sequencing and the lack of reliance on complete assembled genomes might lower the barrier to entry for long-read 16S rRNA gene sequencing as a clinical diagnostic tool. Further, the cost-effectiveness of 16S rRNA gene sequencing can be optimized in the clinic via intelligent design of diagnostic reflex testing^50^. Large datasets of 16S rRNA ASVs from various microbes are thus an attractive choice for developing possible specific microbiome-based biomarkers for human diseases where the role of the microbiome has already been well established, such as cancer^51–53^ and diabetes mellitus^54–56^.

Multiple commercially available tests exist for diagnosing BV in symptomatic women with no recommendations for those who are asymptomatic^39^. Amsel’s criteria and Nugent score are mainstays of BV diagnosis, but may vary in clinical settings due to their subjective nature and technician dependency^57^. Multiplex PCR tests offer stronger sensitivity and specificity at greater expense than Nugent score and Amsel’s criteria, but differ in the microbes that they target^39^. This variability in BV diagnostics poses a challenge for uncovering novel BV biomarkers, which can be overcome using sequencing-based technologies. In this study we confirmed the robustness of 16S rRNA gene sequencing of BV tested remnant vaginal swabs (Labcorp NuSwab® test). However, further work is needed to identify BV microbial and metabolic biomarkers from unconventional clinical manifestations of BV, such as from women who suffer from recurrent BV or from those who present with symptoms but are PCR negative. Large prospective cohort studies of individuals afflicted with recurrent BV symptoms utilizing clinically available PCR testing and 16S rRNA gene-based microbiome profiling (and potentially metagenomic sequencing as well) will be invaluable in helping to close these gaps in the care and treatment of BV.

In this study we profiled the vaginal microbiomes of remnant vaginal swabs clinically tested using the Labcorp NuSwab® BV PCR test. Our findings support the accuracy of NuSwab® in identifying BV and provide valuable insights for advancing the diagnostic and treatment options available to patients. These results highlight the potential for 16S rRNA gene profiling in the diagnosis of BV.

## Materials and Methods

### Sample collection, 16S V3-V4 rRNA gene sequencing, and shotgun metagenomic sequencing

75 clinician-collected remnant NuSwab® samples underwent DNA extraction using the ZymoBIOMICS Magbead DNA isolation kit. The 16S rRNA gene V3-V4 hypervariable region was amplified using the 341f (5′-CCTACGGGNGGCWGCAG-3′) and 805r (5′-GACTACHVGGGTATCTAATCC-3′) primers^58^. Library preparation was performed using KAPA HyperPrep and KAPA Library amplification kits, and the resulting libraries were sequenced on an Illumina MiSeq® platform using 300bp paired-end reads. A separate set of 54 remnant vaginal swabs were processed through the aforementioned sequencing workflow and through a shotgun metagenomic assay comprised of total microbial DNA extraction via the ZymoBIOMICS Magbead DNA isolation kit, library preparation using the KAPA HyperPlus Library Preparation Kit, and sequencing on the Illumina NextSeq2000® platform using 150bp paired-end reads.

### DADA2 processing and taxonomic analysis

16S rRNA gene V3-V4 sequencing reads were processed using R v4.1.1^59^ by first reorienting reads to the same strand and then trimming using the DADA2 v1.22.0^60^ filterAndTrim function with default parameters and trimRight=c(68, 52) (optimally chosen using in-house R function). Quality filtered trimmed reads were then processed through the standard DADA2^60^ process, denoising and merging reads to generate Amplicon Sequence Variants (ASVs). ASVs were then filtered by removing singletons and those with lengths shorter than 350bp. The resulting ASVs were taxonomically classified using the SILVA v138 database^61^ and DADA2 implemented RDP classifier^62^ with a bootstrap confidence threshold of 80. ASVs without at least a Phylum rank classification were removed from analysis.

Additional ASV speciation was performed by aligning ASVs with a genus rank but no species rank classification directly to the SILVA v138 database^61^ using VSEARCH^63^ with parameters “--id 0.99 --strand both --maxgaps 0 --minwordmatches 0 --maxaccepts 0”. Aligned ASVs were assigned to a species if all optimal alignments matched to a single species. *Lactobacillus*-focused speciation was then performed by 99% clustering of all ASVs classified in the *Lactobacillus* genus using CD-HIT^64^. ASVs found in clusters with ASVs previously taxonomically assigned to only a single *Lactobacillus* species (i.e. via RDP classification or direct alignment), were assigned to the same *Lactobacillus* species. Since the NuSwab® BV assay targets microbes not in the SILVA nomenclature (*BVAB-2)* and with phylotype resolution (*Megasphaera-1*), ASVs with genus rank classification were further taxonomically analyzed by direct alignment using VSEARCH as described above to a custom database of 16S sequences obtained from the Bacterial and Viral Bioinformatics Resource Center (BV-BRC)^65^ and additional *BVAB-2* and *Megasphaera* phylotype reference sequences^66,67^. ASVs with uniquely optimal alignments to *BVAB-1*, *BVAB-2*, *BVAB-3*, *Megasphaera-1*, or *Megasphaera-2* were assigned accordingly at the species rank. A final taxonomically aggregated count matrix was then generated by aggregating ASVs at the lowest assigned taxonomic rank.

### Phylogenetic, diversity, and Community State Type analysis

Phylogenetic analysis was performed by generating a multiple sequence alignment of all ASVs using Clustal Omega v1.2.4^68^ and processing it in R using the phangorn package^69^. A neighboring joining tree was constructed using the dist.ml and NJ functions. Then, a Jukes-Cantor optimized maximum likelihood tree was generated using the pml and optim.pml functions with optNni = T. The ggtree and dendextend R packages^70,71^ were used to visualize the resulting phylogenies.

Sample alpha diversity and Community State Types (CST) were determined using the final taxonomically aggregated count matrix (see section ‘DADA2 processing and taxonomic analysis’). Alpha diversity was calculated using the diversity function in the vegan R package^72^. CSTs were determined using the most abundant taxon detected, following the criteria used by Ravel *et al*^12^ (CST-I: *L. crispatus*, CST-II: *L. gasseri*, CST-III: *L. iners*, CST-IV: diverse communities, CST-V: *L. jensenii*).

### Statistical analysis

The relationship between sample BV status and CST classification was examined using Fisher’s Exact tests using the fisher.exact function in R^59^. Differences in microbiome alpha diversity based on BV status and CST were examined using Wilcoxon rank-sum tests using the wilcox.test function in R^59^.

### Confirmation of 16S rRNA gene-resolved CSTs via shotgun metagenomic sequencing

To validate the 16S-based CST characterization, an additional set of 54 remnant vaginal swabs were sequenced via 16S rRNA gene V3-V4 sequencing as described above and via shotgun metagenomics (MGx). The CST classification obtained from the 16S and MGx were compared to validate the reliability of CST types from the 16S pipeline. The MGx reads were processed using Illumina bcl2fastq v2.2 and then filtered using BBMAP v38.98^73^ to ensure high-quality reads with a minimal length of 100 bp. Taxonomic assignments were performed using Kraken v2.1.2^74^, followed by Bracken v2.7^75^ using the pre-built standard plus Refseq protozoa and Fungi (PlusPF) database. CSTs were determined as described above in section ‘Phylogenetic, diversity, and Community State Type analysis’ using the most abundant species detected from the Bracken read re-assignment output (MGx) and the most abundant taxon from the 16S data processing.

### Differential abundance analysis of microbes

The differential abundance analysis of microbes between BV-POS and BV-NEG samples was performed by ANCOM-BC R package^76^. The three BV-indeterminant samples were excluded. Significantly enriched taxa were identified with FDR < 0.05 and L2FC >= 1. Significantly depleted taxa were identified with FDR < 0.05 and L2FC <= −1. The remaining taxa were identified as neutral (FDR >= 0.05 or –1 < L2FC < 1).

### Modularized co-occurrence network analysis

BV-POS and BV-NEG samples were processed using C3NA^38^ to perform modularized co-occurrence network analysis. To account for the cross-taxonomy design and the unique microbial compositional structure, all cross-taxonomy correlations were obtained using Sparse Correlation for Compositional data (SparCC)^77^. Then, the optimal number of taxa clusters was separately determined for BV-POS and BV-NEG samples using a consensus approach implemented in C3NA^38^.

### Prediction and differential abundance analysis of metabolic pathways

Two CSTs with insufficient samples, CST-II with three samples and CST-IV with one sample, were excluded in the metabolic pathway prediction. One outlier BV-NEG sample in CST-IV (NS43) and all three BV-indeterminant samples were excluded. The remaining 67 samples from three major CSTs, CST-I, CST-III, and CST-IV, were used to perform metabolic pathway prediction. The MetaCyc^30^ metabolic pathways of microbes in samples were predicted using their 16S rRNA gene sequences using PICRUSt2^31^ run with default options. Then, ALDEx2^78^ was run with default options to identify pathways at differential relative abundances between different groups of samples. The differential abundance analysis of metabolic pathways was performed on four pairs of groups of samples, including BV-NEG samples within CST-I and BV-POS samples within CST-IV, BV-NEG samples within CST-III vs BV-POS samples within CST-IV, BV-NEG samples within CST-I vs BV-POS samples within CST-IV, BV-NEG samples within CST-I vs BV-NEG samples within CST-III, and BV-NEG samples within CST-IV vs BV-POS samples within CST-IV. Significantly differentially abundant features among different groups of samples were identified using an FDR < 0.05. Differentially abundant pathways were visualized using their centered log-ratio transformed scores and the pheatmap R package^79^. Any pathways described as “superpathways” or “engineered” were excluded from the heatmap.

## Author Contributions

Manuscript conceptualization and methodology – H.N.B., Q.Z., K.S., and L.K.I. Bioinformatics analysis and data processing – H.N.B., Q.Z., K.S., and L.W. 16S rRNA gene sequencing and shotgun metagenomic sequencing – E.A.D. and C.R.I. Preparation of manuscript figures and tables – H.N.B., Q.Z., and K.S. Original manuscript draft preparation – H.N.B and Q.Z. Manuscript review and editing – all authors. Approval of final manuscript – all authors.

## Data Availability

Count matrices, sample metadata, ASV taxonomy and sequences, PICRUSt2 predicted pathway abundances, ANCOM-BC microbial differential abundance results, and ALDEx2 pathway differential abundance results are provided in the **Supplementary Data**.

